# AIM-SNPtag: a computationally efficient approach for developing ancestry-informative SNP panels

**DOI:** 10.1101/427757

**Authors:** Shilei Zhao, Cheng-Min Shi, Liang Ma, Qi Liu, Yongming Liu, Fuquan Wu, Lianjiang Chi, Hua Chen

## Abstract

Inferring an individual’s ancestry or group membership using a small set of highly informative genetic markers is very useful in forensic and medical genetics. However, given the huge amount of SNP data available from a diverse of populations, it is challenging to develop informative panels by exhaustively searching for all possible SNP combination. In this study, we formulate it as an algorithm problem of selecting an optimal set of SNPs that maximizes the inference accuracy while minimizes the set size. Built on this conception, we develop a computational approach that is capable of constructing ancestry informative panels from multi-population genome-wide SNP data efficiently. We evaluate the performance of the method by comparing the panel size and membership inference accuracy of the constructed SNP panels to panels selected through empirical procedures in former studies. For the membership inference of population groups including Asian, European, African, East Asian and Southeast Asian, a 36-SNP panel developed by our approach has an overall accuracy of 99.07%, and a 21-SNP subset of the panel has an overall accuracy of 95.36%. In comparison, the existing panel requires 74 SNPs to achieve an accuracy of 94.14% on the same set of population groups. We further apply the method to four subpopulations within Europe (Finnish, British, Spain and Italia); a 175-SNP panel can discriminate individuals of those European subpopulations with an accuracy of 99.36%, of which a 68-SNP subset can achieve an accuracy of 95.07%. We expect our method to be a useful tool for constructing ancestry informative markers in forensic genetics.

## 1. Introduction

One of the goals in forensic analysis is to ascertain an individual’s ancestry or membership of group origin defined *a priori*, and thus to redeploy investigating efforts. In the past few decades, genetic ancestry-informative markers (AIMs) have been developed for a diverse of populations aiming to infer an individual’s continental or biogeographic origins [1–17]. Recent development of genotyping and sequencing technologies has provided rich sources of genetic makers from diverse populations, enabling AIMs to be fully explored at whole-genome level. In fact, informative genetic polymorphisms have been successfully used to predict physical appearance, such as eye color [18], to infer geographical ancestry [1–17, 19, 20], as well as family of origin [21]. Forensic science is entering a new era of “DNA intelligence” [18].

Among the genetic markers, single nucleotide polymorphisms (SNPs) are the most abundant (approximately one SNP per 1000 bases) and easy to genotype [22]. Furthermore, SNPs can “recover information from degraded DNA samples better than short tandem repeats (STRs)”, which also enables their potential applications in DNA-based forensic tests [16, 23]. However, in most cases, the trace DNA materials available only allow typing of a few genetic loci and the volume of DNA required for whole-genome genotyping using arrays is not plausible in a forensic investigation. Thus, it is pivotal to select panels that include a small number of highly informative SNP loci and meet the specific needs of a forensic investigation.

The promise for such a panel-based approach has already been highlighted by extensive human population genetic studies [24–27]. Many studies revealed that a substantial amount of genetic variations is shared among populations, but only a minor proportion of variations are population specific. Those population-specific SNPs are indicative of an individual’s population origin or membership, however most of these SNPs are in low frequencies, prohibiting them from being used as population tags [28–30]. Among the common SNPs shared by different populations, some SNPs that are highly differentiated by allele frequencies (*i.e*. with high *Fst* scores), are promising candidates for forensic applications and draw a lot of attention in current studies [9, 16]. Moreover, because SNPs are in linkage disequilibria and their information on membership is partially redundant, a non-redundant set that includes a selected number of SNPs could retain the most relevant information needed for ancestry inference. Thus it is possible to condense the number of SNPs without sacrificing their discriminating power for membership inference, thereby being applicable for trace DNA materials.

Several studies have explored such a possibility by developing ancestry informative panels from available large-scale genomic data using empirical procedures *e.g*., [9, 16, 17, 31]. An optimal SNP panel should include a minimum set of SNPs, and the combination of these SNPs can guarantee the highest discriminatory accuracy for ancestral inference. In the present study we developed a computational method satisfying such criteria (named AIM-SNPtag). The method is able to explore the genome-wide SNP data very efficiently, and construct group membership or ancestry informative SNP panels. Testing on an exemplar dataset indicated that our method generated panels that performed better at membership inference with higher inference accuracy, and included fewer SNPs than the panel developed through empirical approaches. The capacity of AIM-SNPtag was further demonstrated by applying it to the 1000 Genomes Project data to generate two continent-level and within-continent SNP panels.

## 2. Materials and Methods

### 2.1 Marker selection procedure

We aim to select a subset of SNPs from whole-genome data that are suitable for ancestry or membership inference. An optimal SNP panel should satisfy two criteria simultaneously: (1) maximize the accuracy of predicting the subject’s membership or achieving a pre-chosen accuracy level, and (2) minimize the number of SNPs required for this inference. However, exhaustively searching through all possible subsets of SNPs to identify a panel that satisfies such criteria is not computationally trivial; the computational time required is an exponential function of the total number of SNPs (n) to be selected against, *i.e*. the computational complexity is O(2^n^). For example, to find a subset from 200 SNPs it requires exhaustively searching and evaluating over 2^200^ possible combinations, which is already impractical. Here we elaborate a method that can explore the genome-wide data more efficiently, while select informative SNP panels approximately satisfying the aforementioned criteria. The method implements a recursive algorithm that downscales the computational complexity to O(*n*^2^).

The proposed method is a forward algorithm that includes four successive steps explained below (Fig. 1) and the relevant mathematical and computational details are in the Appendix.

**Fig. 1.**
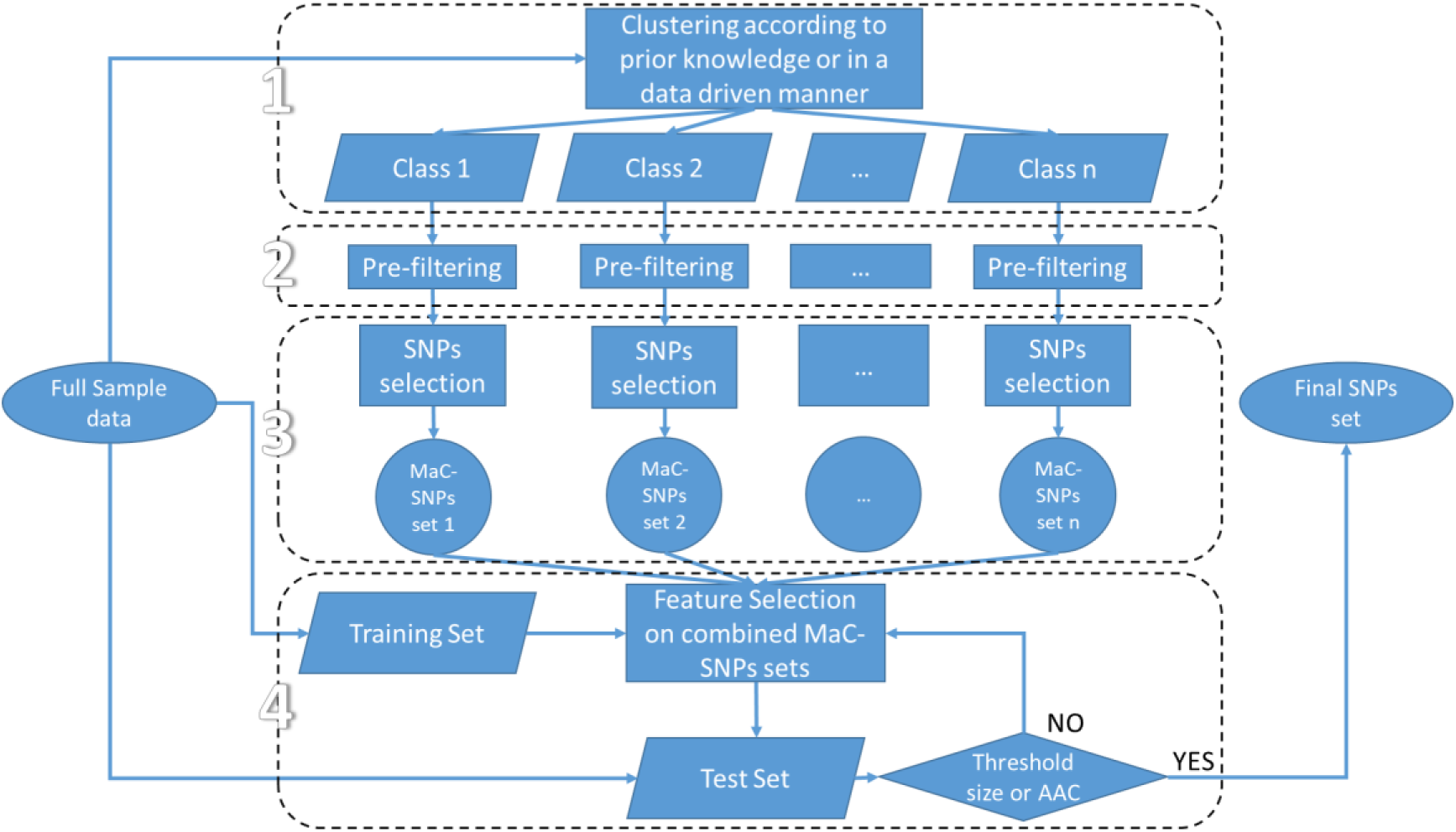
Scheme of AIM-SNPtag for retrieving ancestry or membership informative SNP panel.

Step 1: partitioning. Populations of interest are not equally divergent from each other; in other words, there is inherent hierarchical structure among populations. Consequently, it is difficult to select informative markers that can discriminate populations for all hierarchical levels. Instead of selecting AIMs in a single step simultaneously for all populations, we first partition populations into multiple classes; each class includes two or several populations. The partitioning step can be proceeded with *a prior* knowledge about population structure (such as geographical distribution and/or genetic relatedness) or with a data-driven clustering algorithm such as principal component analysis (PCA). These classes are not necessarily mutually exclusive, instead, they can be nested with each other (see section *Application to genome-wide SNP data* for examples).

Step 2: pre-filtering. When samples include massive number of markers (e.g. >10^6^), a pre-filtering step is necessary to reduce the dimensionality according to the *Fst* values of markers. Since *Fst* measures the degree of differentiation of populations, we first calculate *Fst* of each SNP over populations within the same class. Then we sort the SNPs in a descending order according their *Fst*values. Finally, we retain the top *l* SNPs for each partition class from Step 1. Note that, the number *l* should be much smaller than the total number of SNPs but still reasonably large (say, 20,000), in order to reserve major information that distinguishes populations in the class.

Step 3: selecting. For each class, a feature selection algorithm is proposed to efficiently draw a subset of SNPs (refer to as the maximum classification SNPs, MaC-SNPs) from the pre-filtered SNPs of Step 2. The MaC-SNPs is generated by sequentially adding markers that maximize the accumulative classification ability while panelizing on the maximum normalized mutual information, which account for the effect of linage disequilibrium, over the already added SNPs (see the Appendix for details). As we will integrate the MaC-SNP sets for all classes in step 4, we suggest to apply an even size of 100–200 SNPs for MaC-SNP sets. Indeed, the size is flexible and can be customized according to research demands.

Step 4: integrating and optimizing. The MaC-SNP sets selected from multiple classes are integrated to generate the global AIM-SNP panel. Another wrapper-based feature selection algorithm is applied to generate the global panel which either meets a pre-set threshold of the membership inference accuracy (e.g. 95%) or reaches a pre-given panel size (say, 100 SNPs). The global panel may compose varying numbers of SNPs from each of the MaC-SNP sets (see the Appendix).

The strategies and algorithms summarized above can generate a SNP panel that approximately satisfies the former mentioned criteria, while sharply reducing the computational load, and thus making the method applicable to genome-wide SNP data.

### 2.2 Performance evaluation

We apply the Monte Carlo cross-validation procedure to evaluate the performance of AIM-SNPtag. The full set of labeled samples is partitioned into two subsets, with one subset being the training set which includes 60% of individuals randomly sampled from each labeled population, and the remaining 40% of data is retained as the test set. After constructing the AIM panel with the training set, we apply the Naïve Bayes classifier (NBC, see Appendix) to the test set to evaluate the performance of the AIMs, as NBC is a fast, accurate and reliable classifier. The accuracy, the proportion of correct classifications in test set, 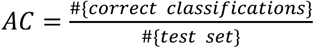 is then calculated. This procedure is repeated for 1000 times. We report the membership inference accuracy as the average accuracy (AAC) over the 1000 repeats.

We further use two pairs of measures to evaluate in more details the performance of classification on every population. They can be derived from a confusion matrix, a 2 × 2 contingency table which records the numbers of four possible outcomes of a binary classification (in terms of the population under investigation), true positive (*TP*), true negative (*TN*) false negative (*FN*), false positive (*FP*). The first pair of statistics are sensitivity 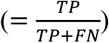 and specificity 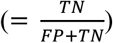, which characterize the distribution of classifications given the true outcomes. The second pair of statistics are positive predictive value 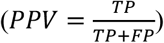 and negative predictive value 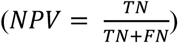, which give the probabilities of the outcomes given the classifications.

The performance of AIM-SNPtag is demonstrated by comparing with existing panels. Given two panels with the same number of SNPs, the higher AAC the better performance the panel has. For a given threshold of accuracy, panels that include fewer SNPs are considered to perform better. As a testing case, we applied AIM-SNPtag to a pre-selected set of 178 SNPs that was also used by [9] to generate a 74-SNP ancestry informative panel (74-AISNP). We generated a panel that discriminate the five major groups, Africans (AFR), Europeans (EUR), East Asians (EA), South Asians (SA), and Southeast Asians (SEA). Then the performance of the AIM-SNPtag selected panel was evaluated using the 1000 Genomes Phase 3 data and compared with the 74-AISNP panel [32].

The informativeness of the panels was also graphically illustrated using the PCA and Bayesian clustering analyses. The PCA analysis was performed using *smartpca* in EIGENSOFT 6.1.4 [33]. The Bayesian method was implemented in STRUCTURE 2.3.4 [34], which assigns individuals into discrete clusters of similar genetic profiles. Ten replicates were run for different fixed number of clusters, *K*, ranging from two to five (*K* = 2~5). For each run, 1×10^6^ iterations were carried out after a burn-in period of 1×10^5^ iterations.

### 2.3 Application to genome-wide SNP data

To further demonstrate its capability, we used AIM-SNPtag to select membership informative SNPs for major world population groups with dense genome-wide SNP data from the 1000 Genomes Project [32]. The five major groups are Africans (AFR, *n* = 108), Europeans (EUR, *n* = 313), East Asians (EA, *n* = 312), South Asians (SA, *n* = 489), and Southeast Asians (SEA, *n* = 192). A total of ~78 million SNPs were collected through whole genome re-sequencing. According to the partition and integration procedure described earlier, we first pre-filtered the SNP pool by retaining 20,000 SNPs for each of the following 3 classes, AFR-EUR-SA-EA/SEA, EUR/SA-EA/SEA and EA-SEA, based on their *F_ST_* scores. From each of the three thinned SNP subsets, AIM-SNPtag exacted a MaC-SNP set with 100 SNPs. These candidate MaC-SNP sets were combined and integrated into global panels that satisfy the thresholds of AAC ≥ 95% and ≥ 99%, respectively.

We also used AIM-SNPtag to select SNP panels that discriminate closely-related subpopulations within Europe. Four local subpopulations, Finnish (FIN, *n* = 99), British (GBR, *n* = 91), Spain (IBS, *n* = 107) and Italia (TSI, *n* = 107), in the 1000 Genomes Project data were analyzed. From the pre-filtered 20,000 SNPs for each of the following 3 classes, FIN-GBR-IBS/TSI, GBR-IBS/TSI and IBS-TSI, we used AIM-SNPtag to select three MaC-SNP sets with 200 SNPs each. These three candidate MaC-SNP sets were then integrated into global panels with AAC ≥ 95% and ≥ 99%, respectively.

## 3. Results

### 3.1 Performance evaluation

From the pre-selected 178 candidate SNPs of the test data set [9], we used AIM-SNPtag to generate an informative panel that discriminates the five major population groups (AFR, EUR, SA, EA and SEA). The average accuracy (AAC) were plotted as a function of the panel size in Figure 2A. The first 18 SNPs selected by AIM-SNPtag discriminate the five major population groups with an AAC of 94.14% peering that of the AISNP-74 panel (94.14%) developed in [9]. A 29-SNP panel (SNPtag-29 hereafter) can achieve an overall AAC of 95.01%. We further examined its discriminative power over each pairs of the five population groups (Fig. 2B). As shown in Fig. 2B, a higher membership inference AAC (> 99%) is achieved if only continental origins were considered. Discriminating EA and SEA origin appeared a bit challenging: in over 86% cases, individual’s origin of EA or SEA were correctly identified. Individuals of EA were mis-predicted as SEA in 13.04% cases and individuals of SEA were mis-predicted as EA in 13.94% cases (Fig. 2B).

**Fig 2.**
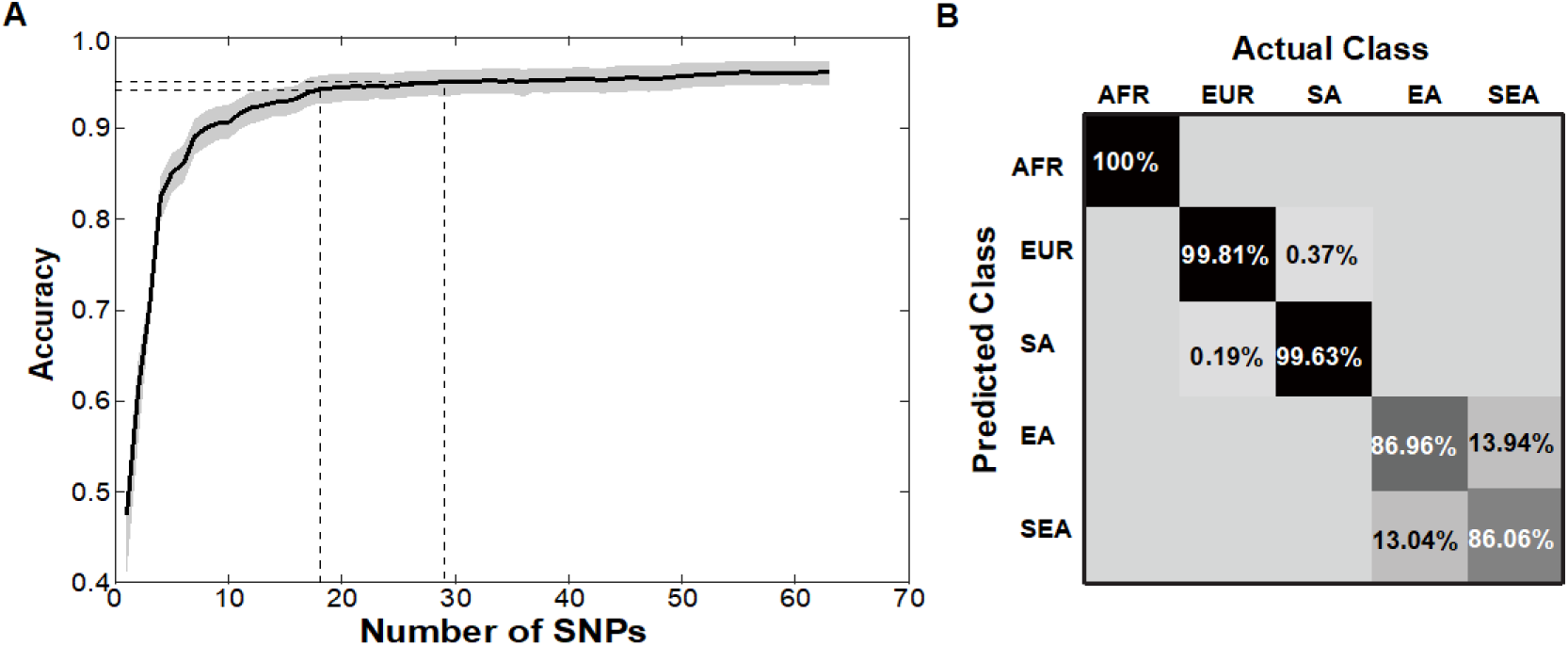
Performance of AIM-SNPtag in retrieving the most informative SNPs from a pool of 178 pre-selected SNPs. (**A**) The plot shows AAC as a function of the panel size. AIM-SNPtag was applied to a pool of 178 SNPs pre-selected in [9] from which the AISNP-74 panel was generated. The horizontal dashed lines indicate the AAC values of 94% and 95% which achieved by the SNPtag-18 panel and SNPtag-29 panel, respectively. The AISNP-74 panel had an AAC value of ~94%. (**B**) The performance of the SNPtag-29 panel in discriminating the five major groups in the 1000 Genomes Project data. The values on the diagonal represent proportion of right inference while those off the diagonal represent proportion of wrong inference.

The performance of the SNPtag-29 panel was further evaluated by four measures and compared to the performance of the AISNP-74 panel [9] on each of the five population groups in Table 1. From Table 1, we can see that SNPtag-29 panel scored higher than the AISNP-74 panel in sensitivity, specificity, *PPV* and *NPV* measurements on all population groups, although it includes less than half number of SNPs of AISNP-74. Results of both the PCA and STRUCTURE analyses also indicate that the SNPtag-29 panel has higher capacity to differentiate the five major population groups than the AISNP-74 panel, especially for the groups within Asia (Fig. 3). In the PCA analysis of the SNPtag-29 data, PC1~PC4 collectively accounted for 56.4% of the total variance. Plot of PC1 (31.1%) vs. PC2 (15.2%) clearly discriminated continental population groups (*i.e*. AFR, EUR, SA and EA/SEA; Fig. 3A), while plot of PC1 vs. PC4 (3.7%) had high power discriminating EA and SEA population groups within Asia (Fig. 3B). Result from STRUCTURE analysis was congruent with PCA analysis where the five population groups were roughly clustered into discrete groups (Fig. 3E). In comparison, both PCA and STRUCTURE analyses of the AISNP-74 panel developed through empirical procedure indicated weaker discriminating power, especially for Asian populations (Fig. 3CDF).We listed details of the 29-SNP panel together with AISNP-74 in Table S1.

**Fig 3.**
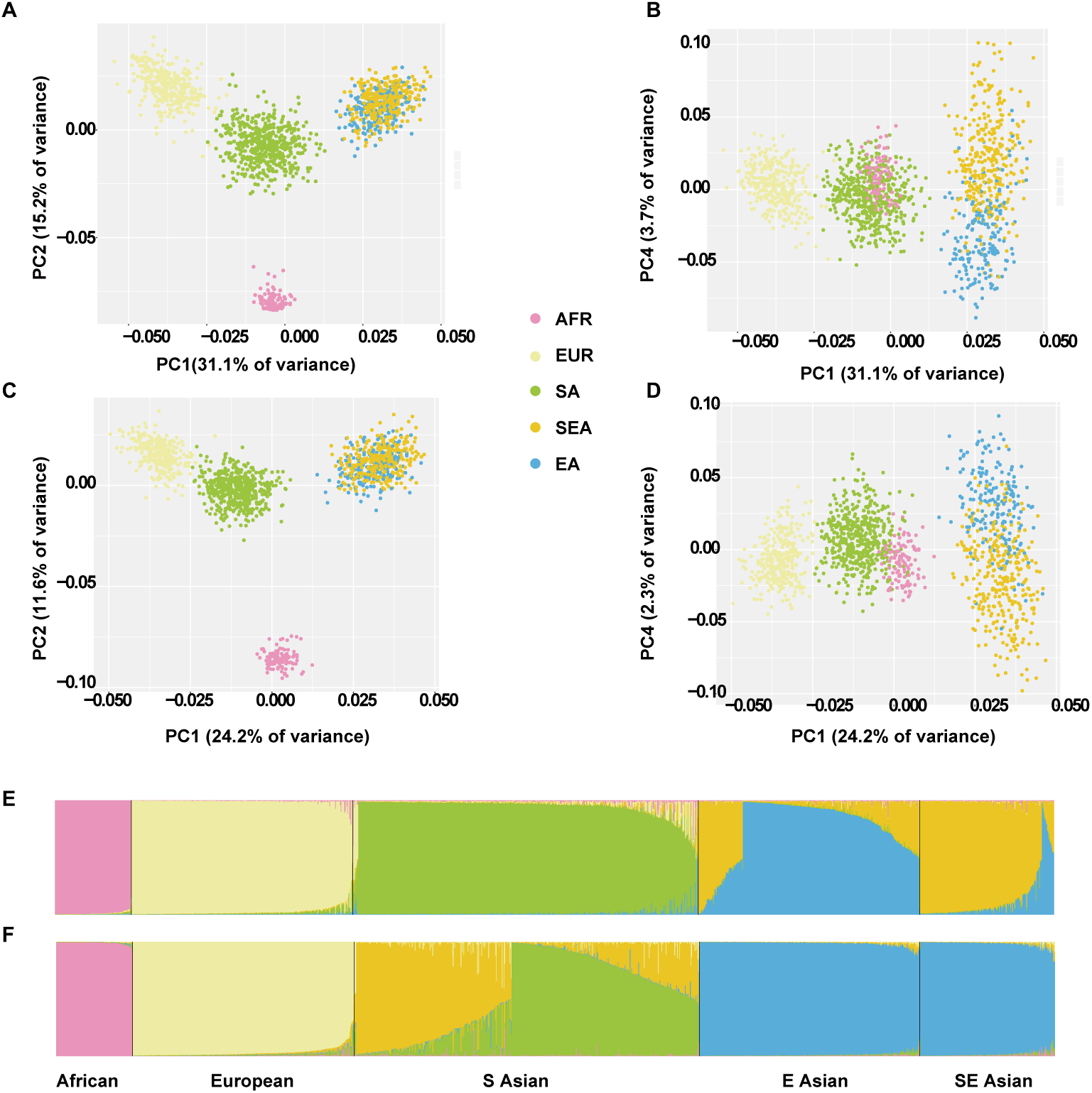
PCA (**A~D**) and STRUCTURE (**E~F**) analyses of individuals from five populations (AFR, EUR, EA, SA, and SEA) using the new SNPtag-29 panel (**AB&E**) the AISNP-74 panel (**CD&F**). Both indicate a better classification performance of the 29-SNP panel.

**Table 1.** Comparison of performance of two SNP panel developed from 178 pre-selected SNP pool on discriminating five major population groups in the 1000 Genome Project data. Values of sensitivity, specificity, PPV and NPV for the 74 AISNP panel (front, regular) and the corresponding values for the SNPtag-29 panel (rear, in bold) are show together.

|  | Sensitivity | Specificity | PPV | NPV |
| --- | --- | --- | --- | --- |
| SAFR | 1.0000/**1.0000** | 1.0000/**1.0000** | 1.0000/**1.0000** | 1.0000/**1.0000** |
| EUR | 0.9996/**0.9981** | 0.9988/**0.9983** | 0.9958/**0.9940** | 0.9999/**0.9995** |
| SA | 0.9973/**0.9961** | 0.9999/**0.9994** | 0.9997/**0.9988** | 0.9986/**0.9980** |
| EA | 0.8448/**0.8713** | 0.9703/**0.9758** | 0.8894/**0.9106** | 0.9567/**0.9640** |
| SEA | 0.8293/**0.8615** | 0.9604/**0.9671** | 0.7668/**0.8051** | 0.9728/**0.9779** |

On the other hand, we should also point out that Li et al [9] includes more minor Asian populations in their data set which are not in the 1000 Genomes Project data. These samples increase the genetic diversity of the EA and SEA groups, and potentially require more AIM markers to discriminate them. However, it is hard to explore its effect on AIM-SNPtag since we don’t have the original data of [9]. Given this fact, we should emphasize more on the discriminating accuracy measures (AAC, sensitivity, specificity, *PPV* and *NPV*) instead of the sizes of panels in the above comparison of both panels when applying to the 1000 Genomes Project data.

### 3.2 New panels for membership inference

We applied AIM-SNPtag to generate ancestry informative SNP panels from genome-wide data from the 1000 Genomes Project [32]. We first generated SNP panels for discriminating the membership for the same five major population groups (AFR, EUR, EA, SA and SEA) as above. Out of the 78 million SNPs, AIM-SNPtag constructed a 21-SNP panel (SNPtag-21) that achieves an overall AAC of 95.36%, and a 36-SNP panel (SNPtag-36) that achieves an overall AAC of 99.07%. The performance of the SNPtag-36 panel on each of the population groups is shown in details in Figure 4A. For all the five population groups, the SNPtag-36 panel can assign individuals to their true populations of origin with an AAC higher than 97%. Only about 2% inference error occurred in distinguishing EA and SEA. The values of four statistics, sensitivity, specificity, *PPV* and *NPV* are high for all population groups, ranging from 0.97 to 1.00 (Table 2).

**Fig 4.**
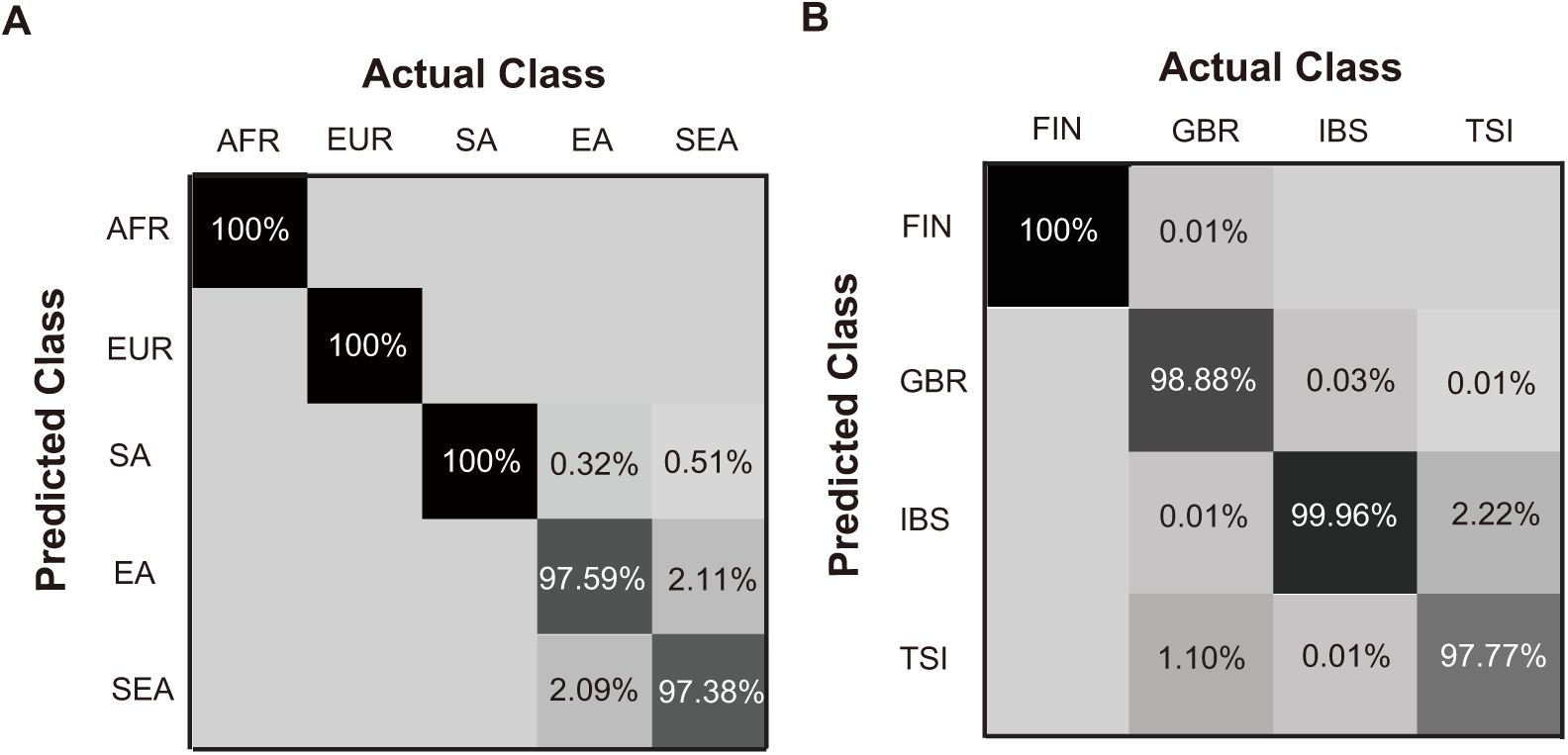
Performance of the SNP panels in discriminating each of the classes. (A) Performance of SNPtag-36 on the five major population groups. (B) Performance of SNPtag-175 on the four subpopulations within Europe. The values on the diagonal represent the proportion of correct inference while those off the diagonal represent the proportion of mis-inference.

**Table 2.** Performance SNP panels developed from genome-wide data sets.

|  | Sensitivity | Specificity | PPV | NPV |
| --- | --- | --- | --- | --- |
| SNPtag-21 (AAC=95%)/SNPtag-36 (AAC=99%) |  |  |  |  |
| AFR | 1.0000/**1.0000** | 0.9998/**1.0000** | 0.9971/**1.0000** | 1.0000/**1.0000** |
| EUR | 0.9992/**1.0000** | 0.9981/**1.0000** | 0.9932/**1.0000** | 0.9998/**1.0000** |
| SA | 0.9935/**1.0000** | 0.9962/**0.9979** | 0.9928/**0.9960** | 0.9966/**1.0000** |
| EA | 0.8303/**0.9759** | 0.9920/**0.9963** | 0.9670/**0.9869** | 0.9538/**0.9932** |
| SEA | 0.9466/**0.9738** | 0.9575/**0.9947** | 0.7786/**0.9663** | 0.9913/**0.9959** |
| SNPtag-68 (AAC=95%)/SNPtag-175(AAC=99%) |  |  |  |  |
| FIN | 1.0000/**1.0000** | 0.9953/**1.0000** | 0.9856/**0.9999** | 1.0000/**1.0000** |
| GBR | 0.9423/**0.9879** | 0.9868/**1.0000** | 0.9541/**1.0000** | 0.9833/**0.9906** |
| IBS | 0.9524/**0.9999** | 0.9587/**0.9934** | 0.8926/**0.9700** | 0.9824/**1.0000** |
| TSI | 0.9081/**0.9813** | 0.9921/**0.9927** | 0.9764/**0.9816** | 0.9677/**0.9926** |

Both PCA analysis and the STRUCTURE analysis indicate that the SNPtag-36 panel reserves high discriminating power for membership of the five population groups (Fig. 5). The first 4 principle components explain nearly half (49.2%) of the variance. PC1 vs. PC2 and PC1 vs. PC3 are highly discriminative of continental population groups while PC2 vs. PC4 distinguish (Fig. 5ABC) regional groups (*i.e*. SA vs. EA vs. SEA) within Asia. All of the five major population groups cluster into nearly discrete blocks with individuals being assigned to their group of origin with high probabilities (Fig. 5D). Compared with the AISNP-74 panel, the SNPtag-36 panel achieves even higher power for discriminating population groups within Asia (Fig. 3F vs. Fig. 6D). The details of these two nested panels were summarized in Table S2. Only three SNPs (rs16891982, rs1426654 and rs17822931) were previously selected in the AISNP-74 panel [9].

**Fig 5.**
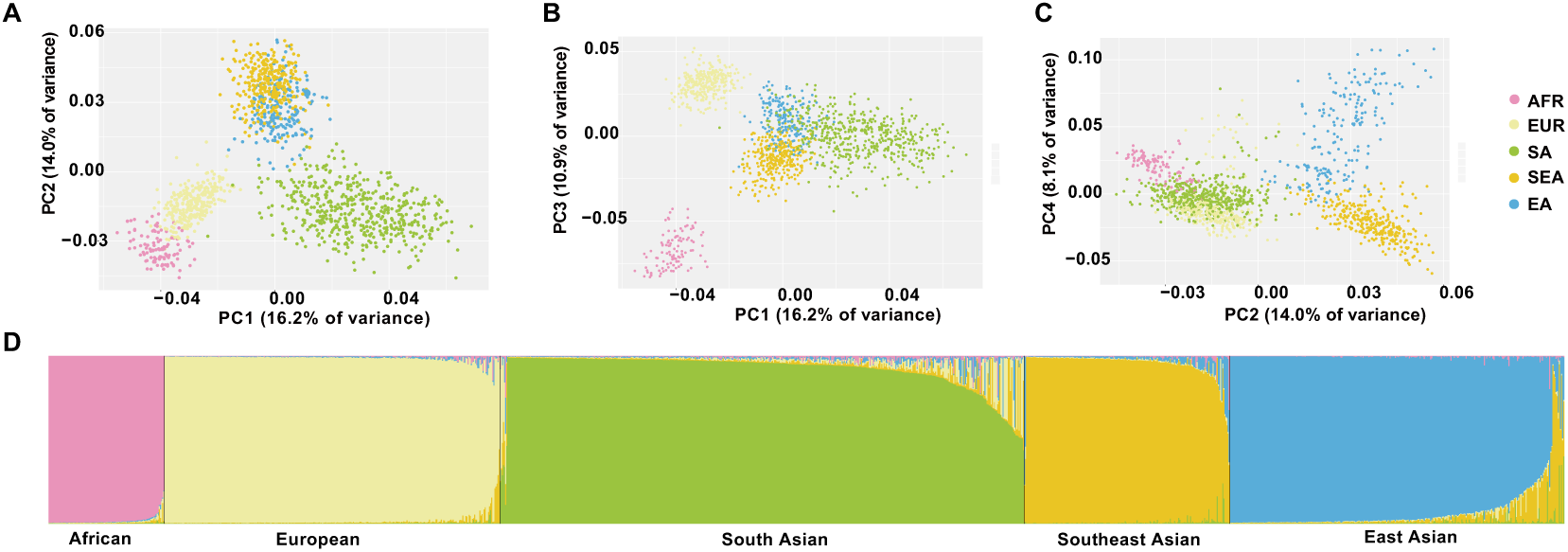
Membership analyses of five reference population groups (AFR, EUR, EA, SA, and SEA) using the SNPtag-36 panel generated by AIM-SNPtag. (**A~C**) Scatter plots of the first four principal components (PC1 vs. PC2, PC1 vs. PC3 and PC2 vs. PC4) discriminate five groups. The first four PCs accounted collectively for 49.2% of the variation in the data. (**D**) STRUCTURE analysis of the individuals.

Similarly, AIM-SNPtag generated SNP panels with high discriminative power in the more challenging case involving four European subpopulations (GBR, FIN, IBS and TSI). AIM-SNPtag generated a 175-SNP panel (SNPtag-175) from the 1000 Genomes Project data, which discriminates the subpopulations with an overall AAC of 99.36%. The first 68 SNPs (SNPtag-68) of the panel achieves an overall AAC of 95.07%. The AAC values for each of the four subpopulation ranges from 97.77% to 100% (Fig. 4B). Small inference error rates occur between subpopulation pairs of TSI vs. IBS (2.22%) and GBR vs. TSI (1.10%). The values of sensitivity, specificity, precision (*PPV*) and *NPV* for the SNPtag-175 panel are all high, ranging from 98.13% to 100% (Table 2). The results of PCA analysis and the STRUCTURE analysis are consistent with the above performance measures (Fig. 6).

**Fig 6.**
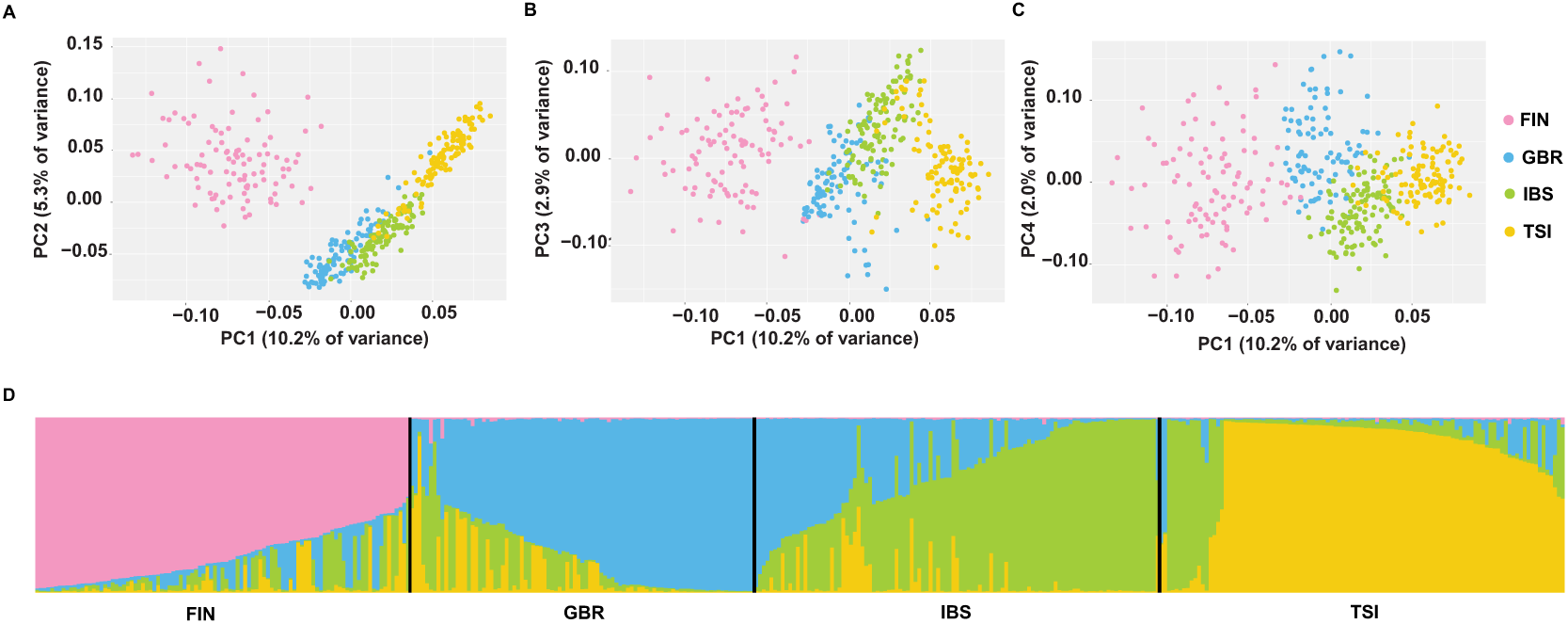
Membership analyses of four European reference subpopulations (GBR, FIN, IBS and TSI) using the SNPtag-175 panel generated by AIM-SNPtag. (A~C) Scatter plots of the first four principal components (PC1 vs. PC2, PC1 vs. PC3 and PC2 vs. PC4) discriminate four groups. (D) STRUCTURE analysis of the individuals.

## 4. DISCUSSION

The deluge of genome-wide SNP data in recent years has enabled us unprecedented opportunities to infer individual’s membership of population groups [19, 20], geographic groups [20] or even linguistic groups [20, 35, 36] with genetic data. However, genotyping a large amount of SNPs is prohibitive in most forensic cases where only trace DNA material is available. Even when ample DNA is available, the intensive computation required for downstream data analysis is time-consuming. Considering these issues, a panel of a relatively small size of highly informative SNPs would be better suited for forensic applications [7, 9, 13].

We developed a computationally very efficient method (AIM-SNPtag) that is applicable to multiple-population genome-wide SNP data and selects the most membership informative SNPs that can be potentially applied to forensic science. Using the proposed algorithm, we successfully downscale the computational complexity of the problem from O(2*^n^*) to O(*n*^2^). AIM-SNPtag generates a panel including a small number of SNPs to achieve a certain degree of accuracy (e.g. 95% or 99%) which could meet the specific purpose.

Two case studies demonstrated that AIM-SNPtag is highly efficient in selecting AIMs for ancestry or membership inference. From millions of SNPs for more than 1,400 individuals, AIM-SNPtag generated a 21-SNP panel (SNPtag-21) that could discriminate the five major population groups with an AAC of 95% and a 36-SNP panel with an AAC of 99% (Fig. 4A). Given the accuracy threshold of 95%, the panels generated by AIM-SNPtag include a small number of markers. In the case of a more challenge situation involving four European subpopulations, GBR, FIN, IBS and TSI, AIM-SNPtag generated a 175-SNP panel (SNPtag-175) with an average accuracy of 99% (Fig. 4B).

We should also point out that in the two case studies, the samples we used to construct AIM panels for the five major population groups (AFR, EUR, EA, SA and SEA) and the European four subpopulations are from the 1000 Genomes Project, and may not be representative of the whole populations. When applying these panels to identify someone from a population not included in the training data, it may cause a bias or mistake. We suggest that in practice AIM-SNPtag should be applied to representative samples of the populations of interest, and users should beware of the AIM panels’ applicable scopes.

In summary, we use the two case studies as illustration examples to demonstrate that AIM-SNPtag can explore the information in the genomic data more sufficiently and efficiently. We believe AIM-SNPtag is a useful tool for SNP panel developments in forensic or medical genetic studies.

## ACKNOWLEDGMENTS

This study is supported by the CAS Key Program (KGFZD-135-16-021), the National Natural Science Foundation of China (91631106 and 31571370), and the “One Hundred Talents Program” of Chinese Academy of Sciences.

## Appendix: Computational algorithms

### Classification ability of a single locus

Denote the full set of loci *L*. Suppose for a given set of populations *K*, we know *a priori P_kl_*(*g*), the frequencies of any genotype *g* ∈ {0,1,2} onlocus *l* ∈ *L* of population *k* ∈ *K*. The differentiation of genotype frequencies from population *j* to population *i* on locus *l* can be described with Kullback-Leibler divergence (KL-divergence) [37] as

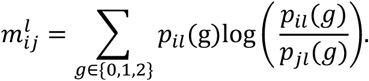

For each locus, the matrix of KL-divergence among all pairs of populations can be written as 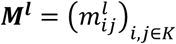. We then define the classification ability of locus *l* as,

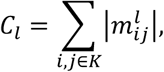

that is, the sum of all elements of matrix ***M^l^***.

### Normalized mutual information between loci

Suppose there are *N* samples. For any loci *l*_1_, *l*_2_ ∈ *L*, denote the genotype frequencies over *N* samples as 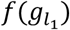 and 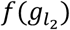, and the joint frequency of genotype pair (haplotype) as 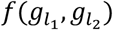, where 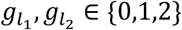. The mutual information (*I*) of loci *l*_1_ and *l*_2_ is then,

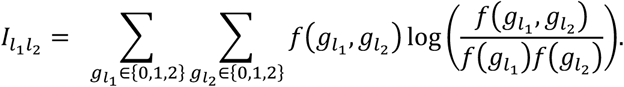

The entropy on any locus *l* ∈ *L* is denoted as

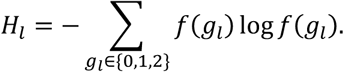

The normalized mutual information [38] between two loci is

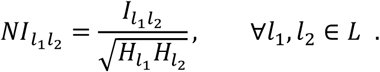

The value of 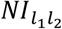 is bounded in [0,1], with 0 indicating complete independence and 1 otherwise.

### Cumulative classification ability of a sequence of loci sets

Let *S_n_ =* {*l*_1_*, l*_2_*,…,l*_3_,…*,l_n_: n ≤ #L;* ∀*i* ∈ {1*, …,n*}*, l_i_* ∈ *L;∀i ≠ j, l_i_ ≠ l_j_*} be a finite sequence of SNPs, where *#L* denote the size of set *L*. For any *i ≤ n*, denote *S_i_ =* {*l*_1_*, l*_2_*,.,l_i_*} the subsequence that constitute of the first *i* items of *S_n_*. We define the cumulative classification ability (CCA) of *S_n_* recursively as follows:

(a). For *S*_1_ = {*l*_1_}*, l*_1_ ∈ *L*, define 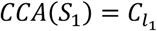, where 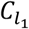 is the classification ability of locus *l*_1_.

(b). For 2 *≤ j ≤n, S_j_ = S_j−_*_1_ ∪ {*l_j_*}, denote,

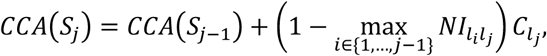

where 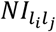 is the normalized mutual information between loci *l_i_* and *l_j_*.

Note that, the calculation of *CCA*(*S_n_*) is dependent on the ordering of the SNPs in sequence *S_n_*.

### Algorithm for selecting the maximum CCA subset

We propose a heuristic algorithm to generate a subset with *u* SNPs that maximize the CCA. We refer to this subset of SNPs as MaC-SNPs (maximum classification SNPs). Our algorithm first generates *#L* sequences of SNPs, each denoted as 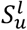, where *l* represents the first item in the sequence is *l* ∈ *L* and *u* indicates the size of the sequence. Each sequence is generated as follows:

(a) For ∀*l* ∈ *L*, 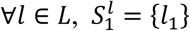, where *l*_1_ *= l*.

(b) For 2 *≤ i ≤ u, 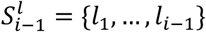, 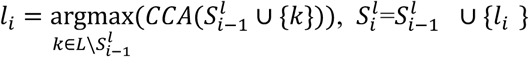*

where 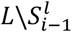 denotes the subset of all SNPs in *L* not in 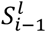. The sequence with maximum *CCA* is selected as the MaC-SNPs, that is, we choose MaC-SNP 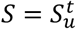, where 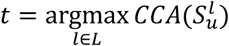.

### Classifier

We adopt the Naïve Bayes classifier (NBC) to test the performance of the MaC-SNP set and infer the population origin of an unlabeled sample. NBC assumes that loci are independent of each other given the targeted label (population). The assumption of conditional independence is rather strong and often violated in practice. Nonetheless, Naive Bayes performs quite well in practice, often comparable to more sophisticated learning methods [39]. For a sample from an unknown population with given genotypes *G =* {*g*_1_*, g*_2_*,.,g_n_*} on the MaC-SNP *S =* {*l*_1_*, l*_2_*, …, l_n_*}, the posterior probability of the sample originated from *k* ∈ *K* is

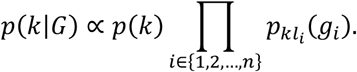

The term *p*(*k*) is the prior probability of population *k*, which can be dropped out if we assume an uniform prior. The 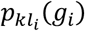 is the genotype frequency of locus *l_i_*∈ *S* in population *k* ∈ *K*, which can be learned by training data with known labels (e.g. samples with known population origin). The label of the sample to be inferred can be obtained by,

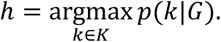

By reducing multiplication to addition the above equation is equivalent to

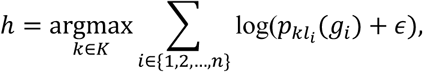

where we add a small positive value *ε* = 10^-4^ to avoid underflow of logarithm calculation.

### Partition and integration

The populations often have hierarchical structure. To generate a SNP panel with discriminative power that comprehensively covers the substructure among available populations, we propose a partition and integration procedure, which extracts the MaC-SNP sets for each of the structured class, and then merges them to generate a final panel of SNPs. The classes can be determined based on *a prior* knowledge, or by clustering methods in a data-driven manner (*i.e*. PCA).

We first extract from each class a candidate MaC-SNP set with a fixed size. The size of the MaC-SNP sets should not be too small. A wrapper-based feature selection algorithm is then applied to obtain the combined global set, which should meet some performance criteria or reach a predefined global set size. Here, we elaborate a greedy forward selection method together with NBC to generate the global SNP panel.

Note, SNPs in each MaC-SNP set are already in an ordered sequence as a result of their generation procedures. The greedy forward selection algorithm for generating the global panel *P* applies as follows. We initialize an empty set *P* and start with the first SNP of each of the MaC-SNP sets. The performances (e.g., average accuracy, AAC, see the main text for definition) of including the first SNP from different MaC-SNP sets to *P* are compared. The SNP that gains the highest AAC is chosen to be a new marker in *P* and it is removed from the corresponding MaC-SNP set. This procedure is repeated until a given threshold of global panel size or AAC is met. The final *P* is thus the global SNP panel.

---

**Table S1.**
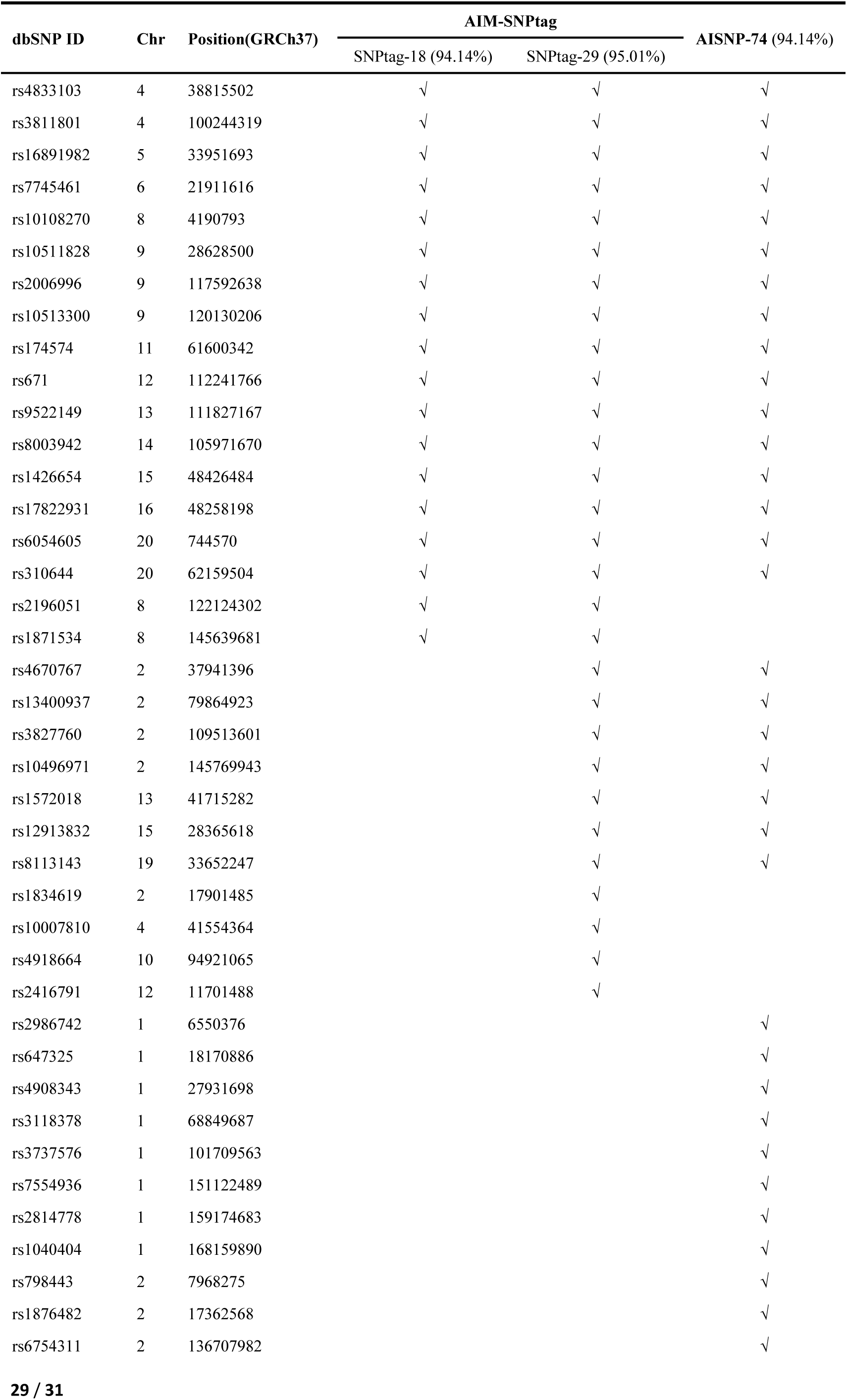
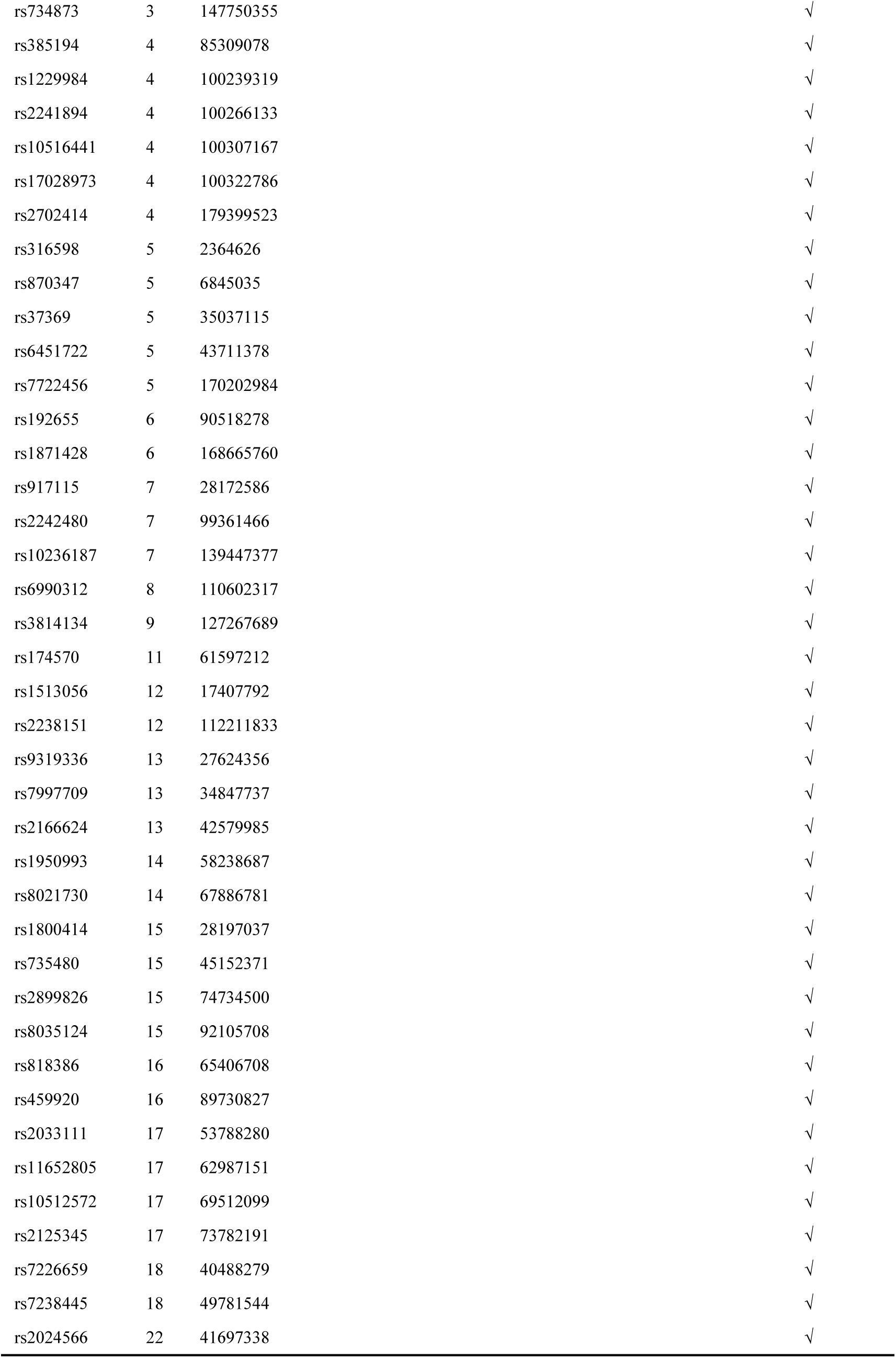
Details of SNP panels selected by AIM-SNPtag from 178 pre-selected SNPs and the AISNP panel by [9].

**Table S2.** Details of two SNP panels selected by AIM-SNPtag from the 1000 Genomes Project data set that discriminate five major population groups. The numbers in parentheses are the membership inference accuracies of the two panels.

| dbSNP ID | Chr | Position<br>(GRCh37) | SNPtag-21<br>(95.36%) | SNPtag-36<br>(99.07%) | Frequencies |  |  |  |  |
| --- | --- | --- | --- | --- | --- | --- | --- | --- | --- |
|  |  |  |  |  | AFR | EUR | SA | EA | SEA |
| rs111824286 | 1 | 604299 | √ | √ | 0.0787 | 0.0272 | 0.0082 | 0.1186 | 0.0000 |
| rs4366944 | 2 | 97628047 | √ | √ | 0.6852 | 0.8962 | 0.3221 | 0.4487 | 0.2812 |
| rs149768401 | 3 | 100365528 | √ | √ | 0.2083 | 0.0815 | 0.0000 | 0.2837 | 0.0078 |
| rs16891982 | 5 | 33951693 | √ | √ | 0.0000 | 0.9201 | 0.0593 | 0.0064 | 0.0052 |
| rs3857620 | 6 | 57496076 | √ | √ | 0.0000 | 0.0000 | 0.4611 | 0.0256 | 0.0052 |
| rs555368174 | 6 | 82912256 | √ | √ | 0.0093 | 0.0208 | 0.0031 | 0.0032 | 0.1615 |
| rs4467891 | 7 | 50222195 | √ | √ | 0.3843 | 0.0016 | 0.3875 | 0.0465 | 0.1328 |
| rs12003714 | 9 | 21932366 | √ | √ | 0.8611 | 0.0080 | 0.1176 | 0.0016 | 0.0182 |
| rs10817902 | 9 | 119309609 | √ | √ | 0.0000 | 0.0000 | 0.3466 | 0.2516 | 0.2161 |
| rs72643557 | 11 | 61579427 | √ | √ | 0.0000 | 0.0032 | 0.0235 | 0.5721 | 0.1719 |
| rs538254210 | 14 | 96938945 | √ | √ | 0.0000 | 0.0288 | 0.0061 | 0.0000 | 0.1901 |
| rs11624491 | 14 | 106188045 | √ | √ | 0.6343 | 0.3163 | 0.5460 | 0.6234 | 0.1328 |
| rs1426654 | 15 | 48426484 | √ | √ | 0.0139 | 0.9984 | 0.6851 | 0.0176 | 0.0026 |
| rs118077539 | 16 | 3281045 | √ | √ | 0.0000 | 0.0000 | 0.3190 | 0.0272 | 0.0339 |
| rs200458768 | 16 | 30998152 | √ | √ | 0.0000 | 0.0000 | 0.5051 | 0.0000 | 0.0052 |
| rs368738705 | 16 | 33921593 | √ | √ | 0.0000 | 0.0000 | 0.4049 | 0.0000 | 0.0000 |
| rs367953206 | 16 | 48221771 | √ | √ | 0.0000 | 0.0000 | 0.3650 | 0.0032 | 0.0026 |
| rs434124 | 19 | 54809336 | √ | √ | 0.6157 | 0.1454 | 0.2444 | 0.7869 | 0.2839 |
| rs2208074 | 20 | 53253867 | √ | √ | 1.0000 | 0.4281 | 0.1125 | 0.0000 | 0.0052 |
| rs187760729 | 20 | 62157764 | √ | √ | 0.0000 | 0.0000 | 0.3722 | 0.0032 | 0.0182 |
| rs61703312 | 8 | 116829577 | √ | √ | 0.9352 | 0.0160 | 0.1227 | 0.0609 | 0.0807 |
| rs561510556 | 1 | 12387655 |  | √ | 0.0046 | 0.0383 | 0.0143 | 0.0112 | 0.2422 |
| rs11204850 | 1 | 151540302 |  | √ | 0.0046 | 0.0000 | 0.2955 | 0.2788 | 0.3177 |
| rs371529719 | 2 | 97897283 |  | √ | 0.0000 | 0.0000 | 0.0133 | 0.0144 | 0.1927 |
| rs546642722 | 4 | 17813761 |  | √ | 0.0000 | 0.0320 | 0.0072 | 0.0064 | 0.2474 |
| rs56357984 | 4 | 38806462 |  | √ | 0.0000 | 0.2748 | 0.1575 | 0.4087 | 0.1094 |
| rs185365488 | 4 | 101002868 |  | √ | 0.0000 | 0.0000 | 0.0092 | 0.0369 | 0.2656 |
| rs201599166 | 4 | 114275541 |  | √ | 0.1620 | 0.2508 | 0.3262 | 0.1571 | 0.4010 |
| rs541669109 | 5 | 44671775 |  | √ | 0.0602 | 0.0351 | 0.0000 | 0.0978 | 0.0000 |
| rs172447 | 9 | 4859106 |  | √ | 0.9630 | 0.0367 | 0.1227 | 0.0401 | 0.0287 |
| rs2001786 | 11 | 128782590 |  | √ | 0.0185 | 0.0000 | 0.2945 | 0.1939 | 0.2318 |
| rs555658336 | 14 | 104193238 |  | √ | 0.1204 | 0.0447 | 0.0000 | 0.1090 | 0.0000 |
| rs201107042 | 16 | 30182239 |  | √ | 0.0648 | 0.0032 | 0.4049 | 0.0609 | 0.1432 |
| rs17822931 | 16 | 48258198 |  | √ | 0.0000 | 0.1102 | 0.4816 | 0.8974 | 0.5885 |
| rs7197127 | 16 | 71473836 |  | √ | 0.9630 | 0.4217 | 0.1196 | 0.0016 | 0.0052 |
| rs389096 | 19 | 54764322 |  | √ | 0.2037 | 0.2444 | 0.3323 | 0.7676 | 0.3958 |

